# DNA-Functionalized Nanoparticles for Multicolor Cathodoluminescence Imaging

**DOI:** 10.64898/2026.04.07.716901

**Authors:** Jeremy B. Conway, Sohaib Abdul Rehman, Maxim B. Prigozhin

## Abstract

Cathodoluminescence (CL) microscopy has the potential to achieve a key goal in biological imaging: the simultaneous visualization of proteins and cellular ultrastructure. This goal can be attained by tagging proteins of interest with spectrally distinct cathodoluminescent probes for detection in electron microscopy. To this end, lanthanide nanoparticles (LNPs) are promising probe candidates due to their stability under the electron beam and their distinct ion-dependent emission spectra suitable for multiplexed detection. However, the hydrophobic surface chemistry of LNPs limits their use in biological samples and requires surface functionalization compatible with aqueous environments and EM sample preparation protocols. Here, we use a DNA-based ligand exchange strategy that renders cathodoluminescent LNPs hydrophilic and compatible with further functionalization for specific protein labeling. We characterize the CL emission of DNA-functionalized LNPs following aqueous transfer and common EM preparation steps, including osmium tetroxide staining and drying protocols based on hexamethyldisilazane and critical point drying, and show that LNPs retain their CL emission under all tested conditions. Finally, we demonstrate multicolor CL imaging of spectrally distinct, DNA-functionalized LNPs on the surface of mammalian cells, enabling simultaneous visualization of cellular ultrastructure via secondary electrons and LNPs via multiple CL color channels.

## Introduction

Precise localization of proteins within the context of cellular ultrastructure (including membranes, chromatin, and the cytoskeleton) is responsible for virtually all aspects of cellular physiology. Unfortunately, no single microscopy technique is well suited to simultaneously image the cellular ultrastructure and specific proteins. Electron microscopy (EM) enables ultrastructural imaging at nanoscale spatial resolution, but has limited molecular specificity. Conversely, fluorescence microscopy enables targeted molecular localization but cannot intrinsically resolve unlabeled cellular ultrastructure. This molecular and ultrastructural information is typically combined using correlative light and electron microscopy (CLEM), which involves sequential imaging of the same sample with fluorescence and electron microscopes^1^. However, CLEM is complicated by incompatible sample preparation protocols between fluorescence and electron microscopy^2^. Moreover, differences in spatial resolution and labeled targets, together with the different contrast mechanisms of the two modalities, create significant challenges for nanoscale image registration^2^.

Cathodoluminescence (CL) microscopy can overcome the limitations of CLEM techniques by directly generating optical emission via electron beam excitation^3^. Using CL as a contrast mechanism for bioimaging offers several unique advantages. Because free electrons have a much shorter de Broglie wavelength than visible photons, CL imaging can offer nanoscale spatial resolution dependent on the volume over which the electron beam interacts with the sample, i.e., the electron interaction volume, rather than the optical diffraction limit, and is not impacted by optical aberrations. Moreover, in CL imaging both ultrastructural information (via secondary electron (SE) detection) and molecular (via CL) information are acquired simultaneously at each pixel during the same electron beam scan, resulting in intrinsic spatial registration without the need for post-acquisition image alignment **(Fig. 1a)**. In addition, this dual-modality information can be obtained at conventional EM acquisition speeds, avoiding the sequential imaging steps required in CLEM. Finally, because CL originates from a spatially localized electron interaction volume, nanoscale protein localization is less dependent on high photon counts compared to single-molecule localization microscopy, where localization precision scales with the square root of the number of detected photons.

**Figure 1.**
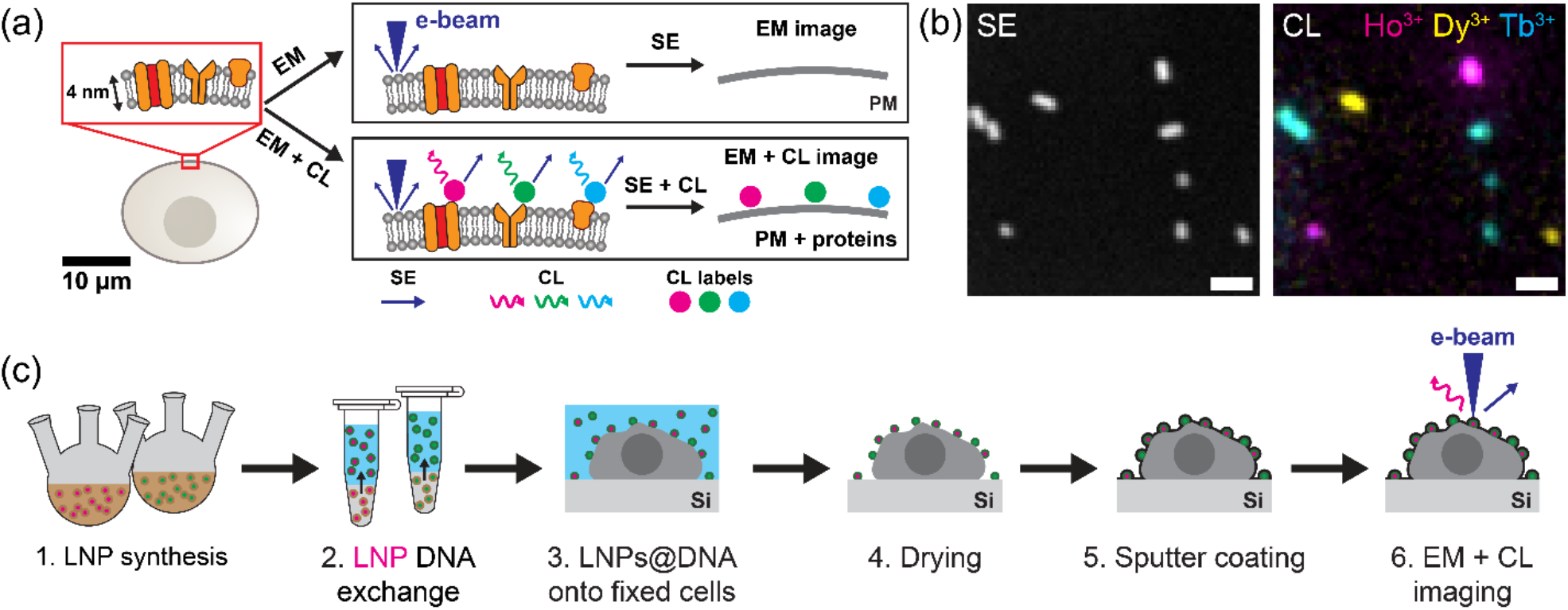
Principle of applying multicolor cathodoluminescent lanthanide nanoparticles (LNPs) to biological electron microscopy. **(a)** Illustration of LNPs as protein labels, allowing simultaneous nanoscale imaging of cellular ultrastructure (such as the plasma membrane (PM)) and specific proteins via EM. **(b**) SE image and corresponding three-color CL image of NaHoF_4_, NaDyF_4_, and NaTbF_4_ LNPs on a Si substrate. **(c)** Illustration of a typical workflow for CL imaging of LNPs in biological samples, from LNP synthesis to sample imaging via EM. To be useful for biological imaging, LNPs must retain their CL properties throughout these steps. Scale bars: (b) 100 nm.

Availability of cathodoluminescent protein labels is central to translating the advantages of CL contrast to bioimaging. To this end, sodium-fluoride-based lanthanide nanoparticles (LNPs)—inorganic nanocrystals that contain ions from the lanthanide series—are promising candidates. They are stable under the electron beam and have narrow spectral features, making them suitable for multiplexing using distinct lanthanide dopants (e.g., Ho^3+^, Dy^3+^, Sm^3+^, Tb^3+^)^4-6^ **(Fig. 1b)**. Furthermore, LNPs generate strong contrast in electron images (refs.^5,6^ and **Fig. 1b**) which enables their precise spatial localization, while CL emission provides spectral identification. This decoupling of detection and classification further reduces reliance on high photon counts for nanoscale protein detection. Previously, we achieved multicolor CL imaging of LNPs on a Si substrate and in biological samples^6^. These results demonstrated the potential of LNPs for single-particle CL imaging. However, the LNPs used in the study were not functionalized for specific protein labeling.

Functionalizing the LNPs for protein labeling is challenging because synthesis methods that provide precise control over LNP size and shape, such as coprecipitation, yield LNPs capped with hydrophobic ligands. These surface ligands preclude their suspension in aqueous solvents. Therefore, the surface chemistry of the LNPs must be altered prior to their application as protein probes. Functionalization methods developed for optically addressable upconversion LNPs^7^ provide a useful starting point for cathodoluminescent LNPs because both types of LNPs have similar crystal lattice structure and surface chemistry. These functionalization techniques generally involve two steps: rendering the surface hydrophilic for colloidal stability in aqueous environments, and introducing functional groups for molecular targeting. Common strategies include silica coating^8,9^, encapsulation with amphiphilic polymers^10,11^ or amphiphilic lipids^12^, oxidation of unsaturated carbon bonds on hydrophobic surface ligands^13^, and direct ligand exchange using hydrophilic ligands^14^.

In this work, we employ DNA ligand exchange in which DNA oligonucleotides replace hydrophobic ligands due to a strong affinity between the LNP surface and phosphate groups of the DNA^15-17^, producing hydrophilic, colloidally stable cathodoluminescent LNPs that can be further functionalized by using common oligonucleotide modifications (e.g., amine groups)^18,19^. Because DNA can be modified prior to coating the LNPs, this approach provides a one-step process that simultaneously achieves ligand exchange and surface functionalization, minimizing particle aggregation that can occur during multi-step surface modifications. We also characterize CL emission from these DNA-functionalized LNPs (LNPs@DNA) after multiple steps commonly used in EM sample preparation, and demonstrate multicolor CL imaging using LNPs@DNA in a biological sample. These results show that after DNA functionalization and EM sample processing **(Fig. 1c)**, LNPs retain their single-particle CL emission, a key requirement for nanoscale CL imaging.

## Results and Discussion

### Coating cathodoluminescent LNPs with DNA

Cathodoluminescent LNPs were synthesized via coprecipitation in a mixture of oleic acid and octadecene, which allowed precise control over nanoparticle nucleation and growth, yielding oleic-acid-capped LNPs through metal–oleate complex formation and resulting in monodisperse LNPs suspended in the reaction solvent^20^. It was essential to transfer the LNPs from this organic medium to a hydrocarbon-free environment for CL imaging because excess hydrocarbons could degrade under electron beam irradiation, leading to contamination and signal loss^21^. The hydrophobic oleic acid coating rendered the LNPs dispersible only in nonpolar solvents such as n-hexane, which was suitable for characterizing the morphology and brightness of LNPs because it rapidly evaporated when added to a substrate, yielding isolated LNPs. However, these hydrophobic LNPs could not be used in hydrophilic, biocompatible solvents.

To make LNPs hydrophilic, we surface-functionalized them with 30-thymine (T30) single-stranded DNA (ssDNA) oligonucleotides by sequentially exchanging the solvent from hexane to chloroform, and then to an aqueous DNA solution through vigorous mixing of the two solutions **(Fig. 2a)**. To achieve this ligand exchange, we adapted a protocol originally developed for upconversion nanoparticles^16^. This process coated the LNP surface with ssDNA, likely via coordination between the negatively charged phosphate groups in the DNA and the positively charged ions on the LNP surface^15,17^. After functionalization, a large fraction of LNPs transferred to the aqueous phase, as confirmed by SEM imaging **(Fig. 2b)**, demonstrating that DNA facilitated the transfer of LNPs to the aqueous phase by binding to the LNP surface and increasing the hydrophilicity of LNPs. A small number of LNPs were also observed in the aqueous phase after mixing the two solutions in the absence of DNA, although they were significantly less abundant than when DNA was present **(Fig. 2c)**. We attributed this limited transfer to the dissociation of oleic acid and exposure of hydrophilic ions at the LNP surface. Also, these DNA-free LNPs appeared more clustered than the DNA-functionalized LNPs, suggesting that DNA functionalization reduces LNP aggregation in the aqueous phase.

**Figure 2.**
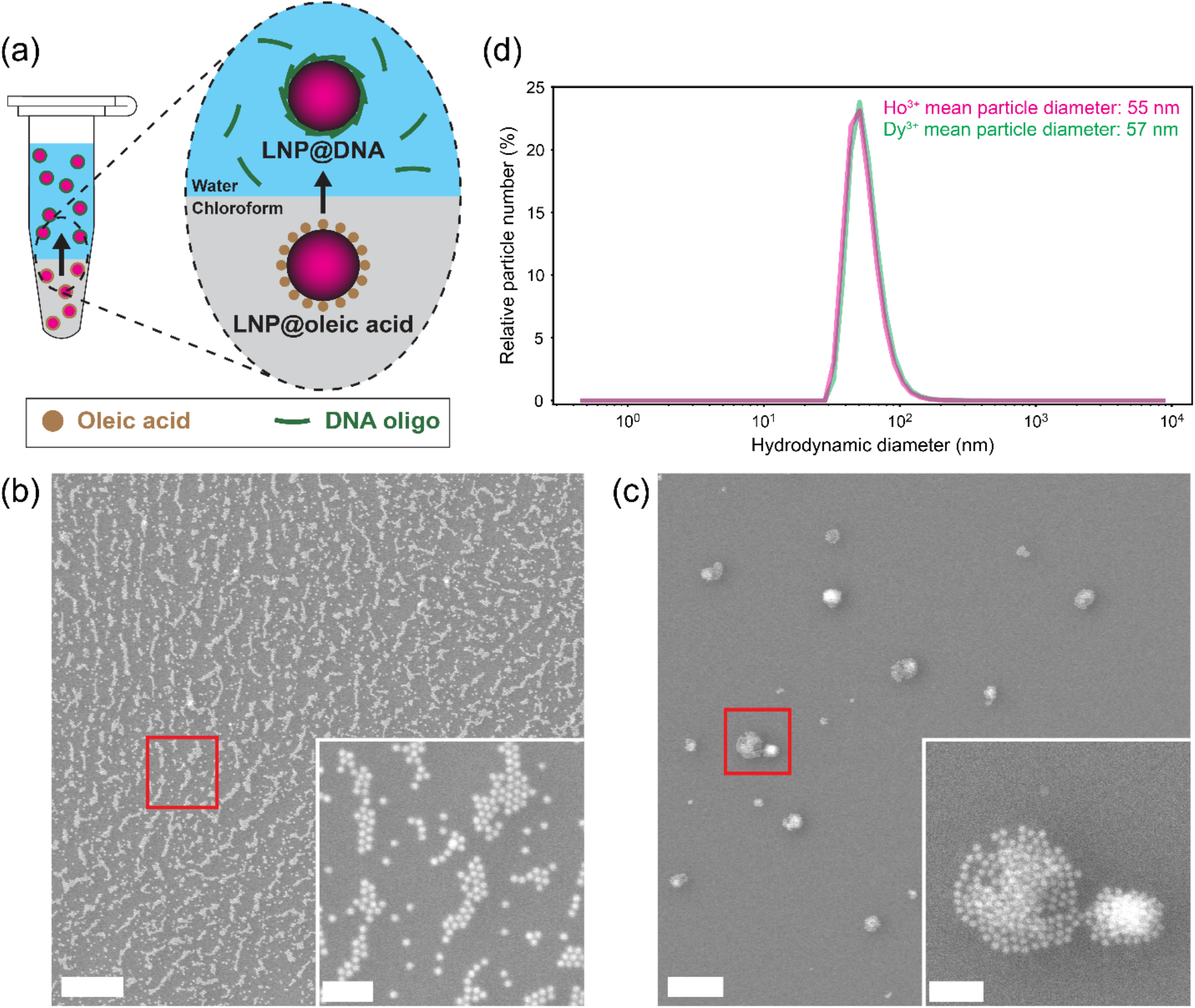
Exchanging hydrophobic LNPs into aqueous solution by surface functionalization with DNA. **(a)** Illustration of LNP functionalization. Initially hydrophobic, oleic-acid-capped LNPs are coated with short DNA oligonucleotides, rendering them hydrophilic. **(b)** SE images of the water layer after performing the exchange protocol with NaHo_0.8_Lu_0.2_F_4_ LNPs and T30 ssDNA. The water layer was drop-cast onto Si and air-dried before imaging. The red square shows the location of the zoomed-in inset. **(c)** SE images of the water layer after performing the exchange protocol with NaHo_0.8_Lu_0.2_F_4_ LNPs but without T30 ssDNA. The water layer was drop-cast onto Si and air-dried before imaging. The red square shows the location of the zoomed-in inset. **(d)** Particle number distributions from dynamic light scattering measurements of NaHo_0.8_Lu_0.2_F_4_ and NaDy_0.8_Lu_0.2_F_4_ LNPs@DNA in water. Mean particle diameters were 54.9 ± 17.0 nm and 57.3 ± 18.7 nm, respectively. Before functionalization with DNA, LNPs were 13.4 ± 1.2 nm and 13.2 ± 3.0 nm in diameter, respectively. Scale bars: (b, c) 500 nm, (b, c insets) 100 nm.

Next, we asked whether the LNPs remained suspended in solution after they were transferred to the aqueous phase. To this end, we performed dynamic light scattering (DLS) measurements on the LNPs 45–60 min after conclusion of the DNA exchange step **(Fig. 2d)**.

This time duration was chosen because it would be sufficient for most labeling strategies, such as those involving self-labeling enzymes (e.g., HaloTag, SNAP-tag) or immunolabeling steps. The DLS analysis showed a single peak centered at 54.9 nm with a standard deviation of 17.0 nm for Ho^3+^-doped LNPs, and a similar peak at 57.3 nm with a standard deviation of 18.7 nm for Dy^3+^-doped LNPs. This apparent increase in hydrodynamic diameter compared to as-synthesized LNP size (13.4 ± 1.2 nm diameter for Ho^3+^-doped LNPs and 13.2 ± 3.0 nm diameter for Dy^3+^-doped LNPs by TEM) and compared to the hydrodynamic diameter of LNPs in hexane (12.0 ± 3.2 nm for Ho^3+^-doped LNPs, Supplementary Fig. 2), as well as the large standard deviation, could be explained by variability in the thickness of the DNA layer (linear length of one T30 oligonucleotide is ≈10 nm) on monodisperse LNPs, or formation of multi-LNP aggregates with an effective mean diameter of ≈55–57 nm. In either case, DLS revealed that LNPs remained suspended in aqueous solution for 60 min. These results showed that, despite their initially hydrophobic surface chemistry post-synthesis, cathodoluminescent LNPs could be rendered hydrophilic using readily available DNA oligonucleotides. We refer to these DNA-functionalized LNPs as LNPs@DNA.

### DNA-coated LNPs retain CL emission following EM sample preparation

Next, we determined the impact of ssDNA coating and aqueous solvent on the CL brightness of LNPs. CL imaging was performed using a custom-modified SEM integrated with an optical detection system. CL was collected using a parabolic mirror, spectrally separated, and focused onto three photomultiplier tubes (PMTs) for multicolor imaging (see Methods and ref.^6^).

We analyzed LNPs containing three different dopants of the form NaHo_0.8_Lu_0.2_F_4_, NaDy_0.8_Lu_0.2_F_4_, and NaTbF_4_. Lu^3+^ ions were co-doped with emitting Ho^3+^ and Dy^3+^ ions to obtain spherical LNPs (Supplementary Fig. 1). This co-doping strategy did not affect CL brightness (Supplementary Fig. 3), consistent with our previous results showing that CL brightness saturates beyond ≈20% dopant concentration due to a balance between the increased dopant population and the higher non-radiative energy losses that occur at higher dopant concentrations^6^. For Tb^3+^ ions, full Tb^3+^ doping was used because we found that co-doping with Lu^3+^ resulted in sub-10-nm LNPs that were too dim for reliable single-particle characterization across different conditions. CL was collected in spectral windows corresponding to the characteristic CL emission peaks of Ho^3+^, Dy^3+^, and Tb^3+^ ions, centered at 647.0 nm, 572.6 nm, and 545.2 nm, with full widths at half maximum (FWHM) of 12.9 nm, 14.3 nm, and 8.5 nm, respectively. We refer to these spectral channels as Ho^3+^, Dy^3+^, and Tb^3+^ channels. Details of the CL imaging setup, acquisition parameters, and data analysis are provided in Materials and Methods and ref.^6^.

**Figure 3.**
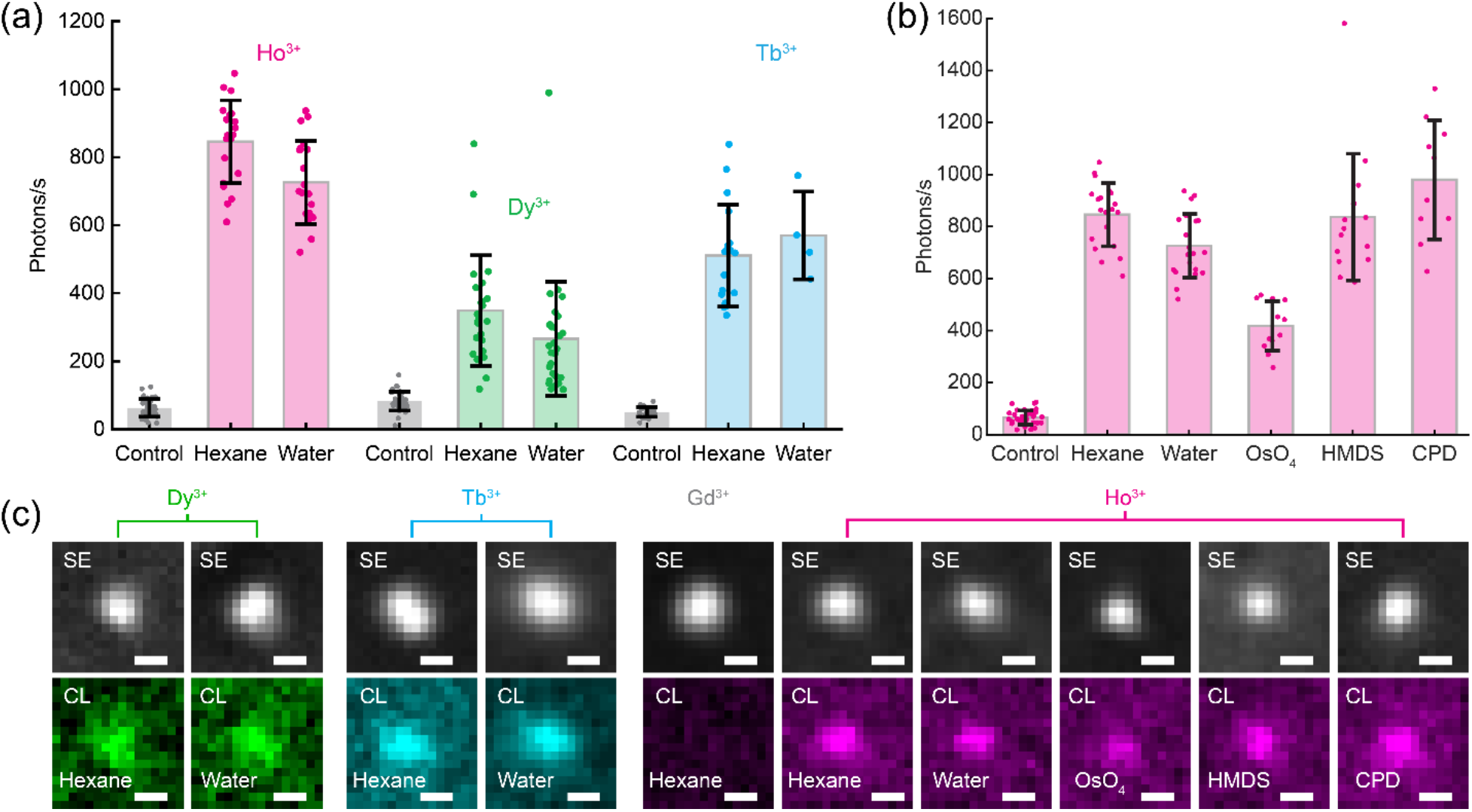
Single-particle CL brightness of LNPs after functionalization and EM sample preparation. **(a)** Single-particle CL brightness of NaHo_0.8_Lu_0.2_F_4_, NaDy_0.8_Lu_0.2_F_4_ (Lu omitted in figure labels for conciseness), and NaTbF_4_ LNPs after DNA functionalization and suspension in water. Corresponding hexane-suspended, oleic-acid-capped LNPs and non-emitting control NaGdF_4_ LNPs are included for comparison. **(b)** Single-particle CL brightness of NaHo_0.8_Lu_0.2_F_4_ LNPs after different treatments typically used for EM sample preparation. After DNA functionalization and suspension in water, LNPs were treated with osmium tetroxide (OsO_4_), hexamethyldisilazane (HMDS), or critical point drying (CPD). Non-emitting control NaGdF_4_ LNPs and hexane-suspended NaHo_0.8_Lu_0.2_F_4_ LNPs are included for comparison. **(c)** Representative SE and CL images of LNPs@DNA used in (a, b). Error bars in (a, b) show the mean and standard deviation of the data. Scale bars: (c) 20 nm.

A comparison of the CL emission from single hydrophobic LNPs prepared in hexane and from LNPs@DNA prepared in water are shown in **Fig. 3a**. The CL brightness of LNPs@DNA was similar to that of post-synthesis hydrophobic LNPs in hexane. After DNA functionalization, we detected on average 726 photons/s from NaHo_0.8_Lu_0.2_F_4_ LNPs, 266 photons/s from NaDy_0.8_Lu_0.2_F_4_ LNPs, and 570 photons/s from NaTbF_4_ LNPs **(Fig. 3a)**. This result shows that the limit on the smallest detectable LNPs@DNA was similar to the limit in hexane, and sub-20-nm LNPs were detectable^6^.

Next, we investigated whether additional steps involved in EM sample preparation protocols affected the luminescence of single LNPs@DNA. These protocols are known to quench the fluorescence of dyes and proteins, primarily due to the use of osmium tetroxide (OsO_4_) for contrast generation in electron microscopy^22-24^, along with subsequent dehydration and drying steps. Thus, the ability of LNPs to retain their luminescence under these conditions is critical for their use as protein probes. We characterized the impact of steps commonly used in SEM sample preparation, specifically OsO_4_ staining and two sample drying methods: hexamethyldisilazane (HMDS) drying and critical point drying (CPD) **(Fig. 3b)**. Across all tested conditions, a CL signal was retained from single LNPs. On average, we detected 418 photons/s from NaHo_0.8_Lu_0.2_F_4_ LNPs@DNA after OsO_4_ treatment, 837 photons/s after HMDS drying, and 979 photons/s after critical point drying. Representative single-particle CL images from each condition are shown in **Fig. 3c**.

A decrease in CL brightness was observed following OsO_4_ treatment. This reduction can be attributed to the strong oxidative nature of OsO_4_ which can alter LNP surface chemistry by crosslinking surface-bound thymines^25^ and residual hydrocarbons, thereby introducing additional nonradiative energy loss pathways. Despite this reduction in brightness, individual LNPs@DNA remained sufficiently bright for reliable single-particle detection and spectral color assignment. Collectively, these results show that LNPs maintained their brightness through multiple EM preparation steps, demonstrating their viability as robust protein labels for CL imaging.

### Imaging LNPs@DNA in a biological sample

Having demonstrated that LNPs@DNA retained their CL properties after standard EM sample preparation steps on a Si substrate, we performed multicolor CL imaging of LNPs@DNA in a biological sample. Specifically, we imaged the LNPs on the surface of mammalian HEK293 cells to assess their emission within the cellular topography. For this proof-of-concept experiment, the LNPs@DNA were not functionalized to target specific proteins. Therefore, to prevent particle loss during washing, we employed a simplified sample preparation protocol: cells were chemically fixed, incubated with an aqueous solution of LNPs@DNA, air-dried, and sputter-coated prior to imaging (see Materials and Methods).

First, we added two spectrally distinct LNPs@DNA (NaHo_0.8_Lu_0.2_F_4_ and NaDy_0.8_Lu_0.2_F_4_) to the surface of cultured mammalian cells **(Fig. 4a)** for CL imaging. For each imaged region, two types of signals were collected: CL images across Ho^3+^ and Dy^3+^ channels, and a SE image to visualize topography of the plasma membrane **(Fig. 4b–d)**. The LNPs exhibited CL emission in either the Ho^3+^ or Dy^3+^ channel **(Fig. 4e)**.

**Figure 4.**
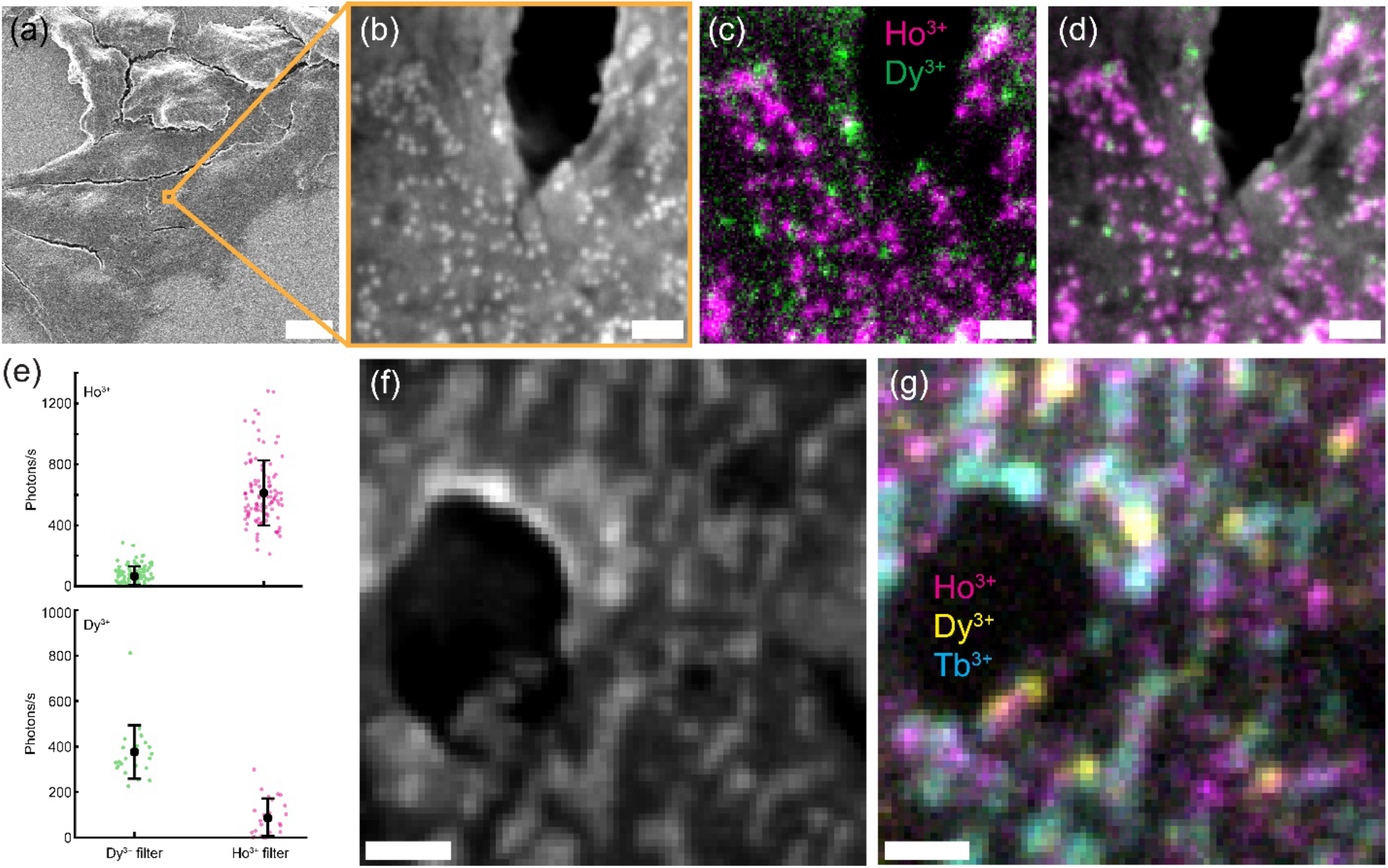
Multicolor CL imaging of LNPs@DNA in biological samples. **(a)** Zoomed-out SE image of HEK293T cells with NaHo_0.8_Lu_0.2_F_4_ and NaDy_0.8_Lu_0.2_F_4_ LNPs@DNA. Cells were chemically fixed and incubated in a water solution of LNPs@DNA, then air-dried and sputter coated with a Pt/Pd mixture for SEM imaging. **(b)** SE image of LNPs@DNA on the surface of a HEK293T cell shown in (a). **(c)** Two-color CL image of the field of view shown in (b). **(d)** Composite SE and CL image of the field of view shown in (b, c). **(e)** Single-particle CL signal from LNPs@DNA on the surface of HEK293T cells, as shown in (b–d). CL signal for NaHo_0.8_Lu_0.2_F_4_ (top) and NaDy_0.8_Lu_0.2_F_4_ (bottom) LNPs@DNA are shown across Ho^3+^ and Dy^3+^ filters. **(f)** SE image of NaHo_0.8_Lu_0.2_F_4_, NaDy_0.8_Lu_0.2_F_4_, and NaTbF_4_ LNPs@DNA on the surface of a HEK293T cell. **(g)** Three-color CL image of the field of view shown in (f). Scale bars: (a) 10 μm, (b–d) 100 nm, (f, g) 200 nm.

Next, we imaged LNPs@DNA of three distinct spectral profiles (NaHo_0.8_Lu_0.2_F_4_, NaDy_0.8_Lu_0.2_F_4_, and NaTbF_4_) on the surface of mammalian cells **(Fig. 4f, g)**. As before, each LNP could be assigned a color based on its emission in the Ho^3+^, Dy^3+^, or Tb^3+^ channel, and topography of the cellular surface was simultaneously visualized in the SE image. The availability of multiple, well-separated emission channels enabled unambiguous identification of individual LNPs even in feature-rich cellular environments, without the need for complex particle classification algorithms that rely solely on SE contrast. These results demonstrate the ability of CL microscopy to simultaneously image cellular ultrastructure and nanoscale probes, and highlight the potential of LNPs as multicolor protein labels for CL imaging.

## Conclusion

In this work, we demonstrated the potential of LNPs as probes for CL imaging, establishing them as a promising platform for visualizing proteins within their ultrastructural context. We showed that cathodoluminescent LNPs could be rendered hydrophilic by coating their surfaces with DNA oligonucleotides, enabling compatibility with aqueous environments. We evaluated the luminescence of LNPs@DNA in water as well as under standard EM sample preparation conditions. In all cases, the LNPs consistently exhibited CL emission. We further applied LNPs@DNA for nanoscale, multicolor CL imaging within a biological sample, where they retained their brightness in the context of cellular ultrastructure. Collectively, these results constitute an advance toward the development of cathodoluminescent LNPs as single-molecule biological labels, allowing for simultaneous molecular identification and ultrastructural imaging within a single high-resolution EM micrograph.

In the future, LNPs@DNA can be functionalized with targeting molecules, such as complementary DNA sequences, aptamers, self-labeling enzymes, antibodies, antibody fragments, and nanobodies, by conjugating them to oligonucleotides. This functionalization would enable the specific targeting of nucleic acids and proteins with multicolor readout. This approach would be analogous to fluorescence in situ hybridization and immunofluorescence or other types of protein labeling in fluorescence microscopy, but would additionally enable simultaneous nanoscale registration of molecular localization with cellular ultrastructure within EM. Such an imaging approach could reveal nanoscale localization of biomolecules in the ultrastructural context of cells and tissues, enabling insights into biological processes such as endocytosis, exocytosis, phagocytosis, autophagy, cell signaling, cell division, cell-cell contact formation, and function of neural circuits.

## Author Contributions

J.B.C., S.A.R., and M.B.P. conceived the project, designed experiments, analyzed the data, and interpreted the results. J.B.C. optimized the nanoparticle synthesis protocol and optimized the preparation of DNA-functionalized nanoparticles. J.B.C. performed DLS and made samples to test different EM sample preparation conditions. J.B.C. and S.A.R. synthesized nanoparticles. S.A.R. developed the CL detection system and implemented the hardware-software integration for simultaneous SEM and CL imaging. S.A.R. performed CL imaging and developed the imaging and data analysis pipelines. J.B.C. and S.A.R. performed TEM imaging. J.B.C. performed cell culture and prepared biological samples for CL imaging. M.B.P. supervised the research. The manuscript was written through contributions of all authors. All authors have given approval to the final version of the manuscript.

## Acknowledgements

The authors thank Debsankar Roy, Simon Merminod, Margaret Doyle, Surabhi Sreenivas, Arvind Srinivasan, Ami Thakrar, Daphne-Eleni Archonta, Jimmy Hsu, Balmiki Kumar, and Karan Malhotra for useful discussions, as well as James MacArthur and Edward R. Soucy for assistance with CL instrumentation.

## Materials and Methods

### Materials for nanoparticle synthesis and sample preparation

Oleic acid (Sigma-Aldrich, Cat# 364525-1L, 90%, technical grade), 1-octadecene (Sigma-Aldrich, Cat# O806-1L, 90%, technical grade), rare-earth chlorides [gadolinium trichloride hexahydrate (Sigma-Aldrich, Cat# 203289-25G, 99.999%), terbium trichloride hexahydrate (Sigma-Aldrich, Cat# 212903-5G, 99.9%), dysprosium trichloride hexahydrate (Sigma-Aldrich, Cat# 289272-25G, 99.9%), holmium trichloride hexahydrate (Sigma-Aldrich, Cat# 289213-5G, 99.9%), lutetium trichloride hexahydrate (Sigma-Aldrich, Cat# 542075-5G, ≥99.99%)], sodium hydroxide (Sigma-Aldrich, Cat# 221465-500G, ACS Reagent, ≥97.0%, pellets), methanol (Sigma-Aldrich, Cat# 34860-1L-R, ≥99.9%), ammonium fluoride (Sigma-Aldrich, Cat# 216011-100G, ACS Reagent, ≥98.0%), n-hexane (Sigma-Aldrich, Cat# HX0302-3, 95%), ethanol (Ethyl alcohol, Sigma-Aldrich, Cat# 459844-1L, 200 proof, ACS Reagent, ≥99.5%), chloroform (Macron, Cat# 4440-04, ACS Reagent), T30 ssDNA (Integrated DNA Technologies, custom oligo order).

### Nanoparticle synthesis

NaTbF_4_ and NaGdF_4_ LNPs were synthesized by the coprecipitation method based on refs.^5,26-28^. Briefly, 0.5 mmol of the appropriate lanthanide chloride hydrate was combined with 3 mL oleic acid and 7.5 mL 1-octadecene. The reaction was placed under vacuum and the temperature was set to 160 °C for 30 min with constant mixing. The solution was cooled to <30 °C. Next, 1.25 mL of 1 M sodium hydroxide in methanol and 5 mL of 0.4 M ammonium fluoride in methanol were combined, then added to the solution. The reaction was mixed at room temperature for 60 min. The temperature was increased to 70–80 °C and maintained for 30 min. Then, for the LNP growth step, the reaction was placed under an argon atmosphere and heated rapidly to 320 °C. This temperature was maintained for 60 min before cooling to <30 °C. For NaHo_0.8_Lu_0.2_F_4_ and NaDy_0.8_Lu_0.2_F_4_ LNPs, the synthesis was performed as described above, except that the 0.5 mmol of lanthanide chloride hydrates was divided into two quantities based on the desired molar ratio of the two lanthanide elements. Also, the 320 °C reaction temperature was maintained for 5–15 min for NaHo_0.8_Lu_0.2_F_4_ LNPs, and 25–30 min for NaDy_0.8_Lu_0.2_F_4_ LNPs, before cooling to <30 °C. LNPs were stored as-synthesized in oleic acid and 1-octadecene at room temperature.

### Hydrophobic nanoparticle sample preparation

To prepare samples of hydrophobic nanoparticles, 0.5 mL of as-synthesized LNPs was mixed with 5 mL ethanol and centrifuged at 5,000 × *g* for 10 min. The pellet was resuspended in 1 mL n-hexane, mixed with 5 mL ethanol, and centrifuged again. This process was repeated for a total of five times. After the fifth wash, the pellet was resuspended in 5 mL n-hexane and the solution was left undisturbed overnight. For further sample preparation, LNP solution was pipetted from the top of the volume to avoid collecting precipitated LNP aggregates. 2 µL of the solution was drop-cast onto a TEM grid, or 1 µL was drop-cast onto an ethanol-washed and plasma-cleaned p-type Si wafer for characterization by TEM or SEM, respectively.

### Hydrophilic DNA-functionalized nanoparticle sample preparation

To prepare samples of hydrophilic nanoparticles, 400 µL n-hexane-suspended LNPs was mixed with 2 mL ethanol and centrifuged at 5,000 × *g* for 10 min. The supernatant was removed then the pellet was air-dried for >15 min and resuspended in 80 µL chloroform. The resulting LNP solution was added to 200 µL 100 µM T30 ssDNA in nuclease-free molecular biology grade water (or, for the DNA-free control experiment, molecular biology grade water without ssDNA).

The water and chloroform phases were shaken together overnight in an additive-free polypropylene tube (to avoid dissolution of additives by chloroform) on a Vortex Genie 2 on the 4/10 RPM setting. The tube was left still for 10 min, then liquid was pipetted from the top of the volume and into another tube. If lower LNP density was desired, this liquid was diluted 2x–10x in molecular biology grade water. 1 uL of the final dilution was drop-cast onto a plasma-cleaned p-type Si wafer and air-dried in a fume hood. The dried sample was imaged via SEM or processed further (see below). If applicable, the remaining volume was used for DLS or was drop-cast onto fixed cells (see below).

### DLS

For DLS with hydrophilic LNPs, as-prepared LNPs@DNA (see above) were diluted 10x in molecular biology grade water, filtered using a 0.22 µm hydrophilic PTFE syringe filter, then particle size was measured using a Malvern Zetasizer Nano ZS. Measurements were taken at 25 °C with the material refractive index set to 1.50, dispersant refractive index set to 1.33 and viscosity set to 0.8872 cP, and using a measurement angle of 173°. DLS measurements were performed 45–60 min after ending the DNA exchange step (see above).

For DLS with hydrophobic LNPs, as-prepared hexane-suspended LNPs (see above) were diluted 10x in n-hexane, then particle size was measured using a Malvern Zetasizer Nano ZS. Measurements were taken at 25 °C with the material refractive index set to 1.50, dispersant refractive index set to 1.38 and viscosity set to 0.31 cP, and using a measurement angle of 173°. DLS measurements were performed immediately after diluting LNPs in n-hexane.

### Treatment of LNPs with OsO_**4**_

After drop-casting LNPs@DNA on a Si wafer and air-drying (see above), the Si wafer was submerged in 1% OsO_4_ in MilliQ water for 10 min, washed three times with MilliQ water, then air-dried in a fume hood. The dried sample was imaged via SEM.

### Treatment of LNPs with HMDS

After drop-casting LNPs@DNA on a Si wafer and air-drying (see above), the Si wafer was submerged in HMDS for 45 min then air-dried in a fume hood. The dried sample was imaged via SEM.

### Treatment of LNPs with CPD

After drop-casting LNPs@DNA on a Si wafer and air-drying (see above), the Si wafer was submerged in ethanol inside a Tousimis 931.GL critical point dryer, and the sample was dried using the instrument’s auto protocol. The dried sample was imaged via SEM.

### CL imaging setup

Imaging was performed as described previously^6^ using a custom-modified ZEISS SUPRA 55VP FESEM, integrated with a CL imaging system. CL was collected by an off-axis, aluminum parabolic mirror with a focal length of 1 mm and a collection solid angle of ≈1.34π steradian. The mirror directed CL out of the vacuum chamber of the SEM, where it was spectrally separated using dichroic mirrors, filtered by band-pass filters, and focused onto photomultiplier tubes (Hamamatsu H7421-40) using 30-mm-focal-length lenses. CL was simultaneously collected over three color channels. The band-pass filters were Chroma ET645/30X (Ho^3+^ color channel), Chroma FF03-575/25 (Dy^3+^ color channel), and Chroma ET550/20X (Tb^3+^ color channel).

### Single-particle CL image acquisition

Single-particle CL imaging was performed with an electron beam energy of 3–4 keV, a beam current of 160–200 pA, and a working distance of 6–7 mm. Both the Zeiss SmartSEM software and custom LabVIEW software were used for image acquisition. A region of interest was imaged with a pixel size of 5 nm (for single-particle brightness characterization) and 7–13 nm (for multicolor imaging). Four images were captured during each scan of the electron beam: one SE image and three CL images in different spectral channels. SE and CL images were smoothed by convolution with a 2D Gaussian function of 0.5–0.7 pixels standard deviation, except for Fig. 1b where the CL image, but not the SE image, was smoothed. To characterize emission properties of single LNPs, 50 frames were captured with a dwell time of 1 ms per pixel per frame to overcome beam induced drift^6^, resulting in an effective beam dwell time of 50 ms per pixel. These conditions translate to a current density of 3–12 pA per nm^2^ and a total electron dose of ≈6.25 × 10^7^ electrons per pixel. For multicolor experiments, images were acquired with a beam dwell time of 0.1–1 ms and an effective beam dwell time of 30–50 ms.

### Drift correction

Drift correction was performed by acquiring multiple sequential frames using a short dwell time (0.1 ms for multicolor experiments and 1 ms for single-particle emission characterization experiments), instead of a single image with a long dwell time. LNPs themselves were used as fiducial markers in SE images. The position of a LNP was determined by fitting a 2D Gaussian function to its SE image. Euclidean distance between the positions of the LNP in successive SE frames was calculated and rounded to the nearest integer. This distance was then used to translate SE and CL images to account for the drift during image acquisition.

### CL emission rate and photons per frame

The CL properties of a LNP were determined by first localizing it in the SE images and then determining its emission properties at the corresponding location in the CL images. The position of the LNP in a SE image was determined by fitting a 2D Gaussian function. The CL image was then fit to a 2D Gaussian function of the form:

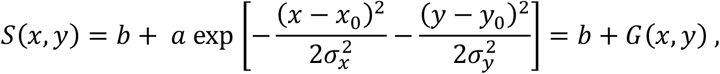

where (*x*_0_, *y*_0_) is the center position of the LNP, *b* is the background, *a* is amplitude, and σ_*x*_ and σ_*y*_ are the standard deviations of the 2D Gaussian function. *G*(*x, y*) is the 2D Gaussian fit to the LNP without the background. In this equation, *x*_0_, *y*_0_, σ_*x*_, and σ_*y*_ were constrained by the 2D Gaussian fit of the SE image. The rate of CL emission was determined as 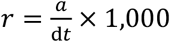, and the total number of photons emitted by the LNP in a frame was calculated by summing over *G*(*x, y*).

### LNP size

The diameter (also referred to as size) of a LNP was determined from the oversampled SE images taken with the ZEISS SmartSEM software (pixel size 0.2–0.6 nm). To determine the diameter of a LNP, a 2D Gaussian function was fit to its SE image. LNP diameter was defined as the FWHM of this fit. FWHM was calculated from the standard deviation of the Gaussian fit as 2.35σ, where σ is the average of the two standard deviations (σ _*x*_ and σ _*y*_) of the 2D Gaussian fit.

To determine LNP diameter from TEM images, a Gaussian blur with a standard deviation of 2 pixels was applied, and the image threshold was adjusted to distinguish LNPs from the background and render a binary image. Using FIJI software, the Fill Holes process was applied to the binary image, then the Analyze Particles function was used to determine each particle’s area with circularity range set to 0.3–1.0 and particle size range set to 40–310 nm^2^ for NaHo_0.8_Lu_0.2_F_4_ LNPs and 40–500 nm^2^ for NaDy_0.8_Lu_0.2_F_4_ LNPs. Finally, each particle’s diameter was calculated as the diameter of a circle with the same area as the particle.

### Biological sample preparation

Cultured cells were prepared for CL imaging via fixation, addition of LNPs@DNA, then air-drying. HEK293T cells were cultured in 50/50 DMEM/F-12 media with 10% fetal bovine serum and 1% penicillin/streptomycin. Cells were dissociated using 0.25% trypsin in PBS, then seeded on plasma-cleaned Si wafers. After adhering to Si overnight, cells were washed twice with PBS and fixed with 4% formaldehyde, 0.2% glutaraldehyde in PBS, pH 7.4 for 15 min. Cells were washed three times with molecular biology grade water, then liquid was removed by pipetting. 20 μL of LNPs@DNA in molecular biology grade water (combined at a 1:1 or 1:1:1 ratio by volume for two- and three-color experiments, respectively) was added onto the cells.

Cells with LNPs@DNA solution were air-dried in a fume hood overnight. Finally, cells were sputter coated with a 2-nm-thick layer of 80:20 Pt:Pd to prevent charging during CL-SEM imaging.

