## Supplemental information for "DNA-Functionalized Nanoparticles for Multicolor Cathodoluminescence Imaging"

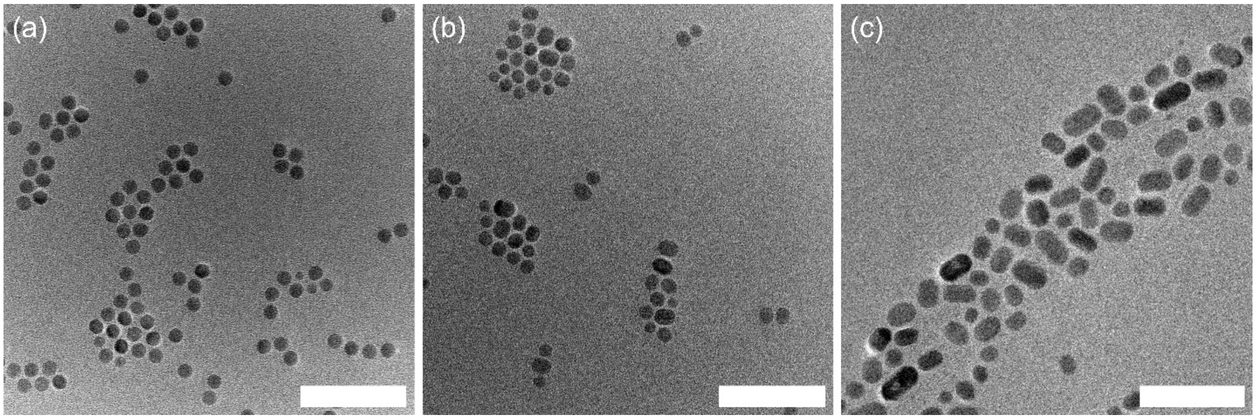


**Supplementary Figure 1: TEM images of LNPs.** TEM characterization of **(a)** $NaHo_{0.8}Lu_{0.2}F_4$, **(b)** $NaDy_{0.8}Lu_{0.2}F_4$, and **(c)** $NaTbF_4$ LNPs. Images were acquired at 200 keV beam energy using a JEOL JEM-F200 S/TEM in TEM mode. Scale bars: 100 nm.

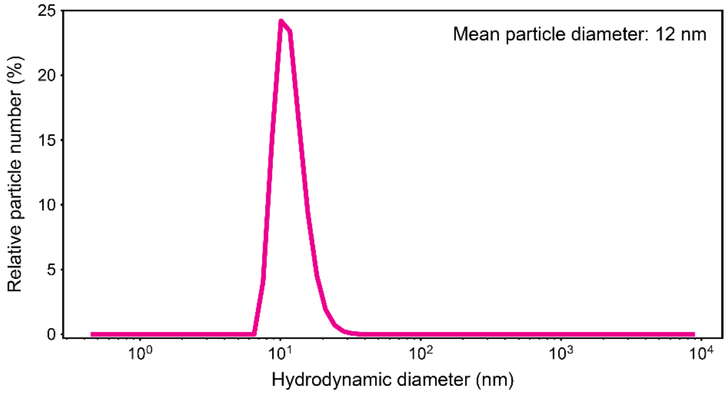


**Supplementary Figure 2: DLS with hexane-suspended LNPs.** Particle number distribution from DLS measurements of $NaHo_{0.8}Lu_{0.2}F_4$ LNPs in n-hexane. Mean particle diameter was 12.0 ± 3.2 nm.

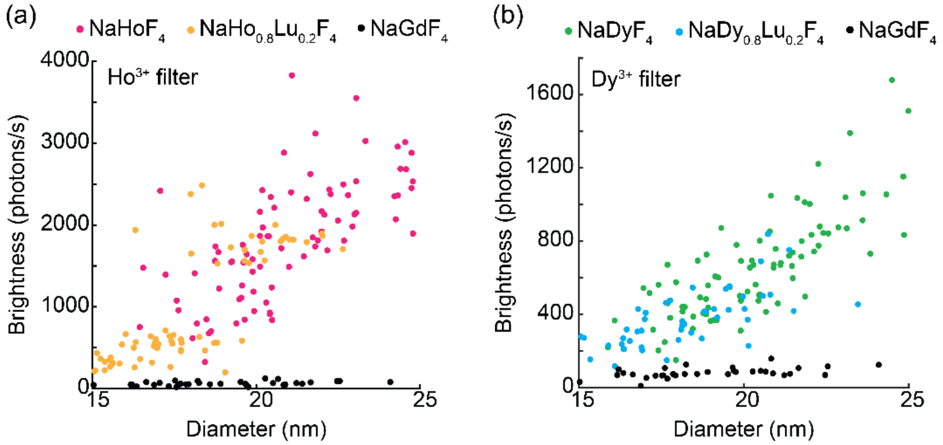


**Supplementary Figure 3: Brightness of LNPs when co-doped with $Lu^{3+}$ ions. (a)** Brightness of $NaHoF_4$ and $NaHo_{0.8}Lu_{0.2}F_4$ LNPs in the $Ho^{3+}$ filter as a function of LNP diameter. **(b)** Brightness of $NaDyF_4$ and $NaDy_{0.8}Lu_{0.2}F_4$ LNPs in the $Dy^{3+}$ filter as a function of LNP diameter. Brightness of control $NaGdF_4$ LNPs is also shown in the respective filters for comparison.
